# Artificial intelligence method to design and fold alpha-helical structural proteins from the primary amino acid sequence

**DOI:** 10.1101/660639

**Authors:** Zhao Qin, Lingfei Wu, Hui Sun, Siyu Huo, Tengfei Ma, Eugene Lim, Pin-Yu Chen, Benedetto Marelli, Markus J. Buehler

## Abstract

The development of rational techniques to discover new proteins for use in variety of applications ranging from agriculture to biotechnology remains an outstanding materials design problem. The key barrier is to design a sequence to fold into a predictable structure to achieve a certain material function. Focused on alpha-helical proteins, we report a Multi-scale Neighborhood-based Neural Network (MNNN) model to learn how a specific amino acid sequence folds into a protein structure. The algorithm predicts the protein structure without using a template or co-evolutional information at a maximum error of 2.1 Å. We find that the prediction accuracy is higher than other models and the prediction consumes less than six orders of magnitude time than *ab initio* folding methods. We demonstrate that MNNN can predict the structure of an unknown protein that agrees with experiments, and our model hence shows a great advantage in the rational design of *de novo* proteins.

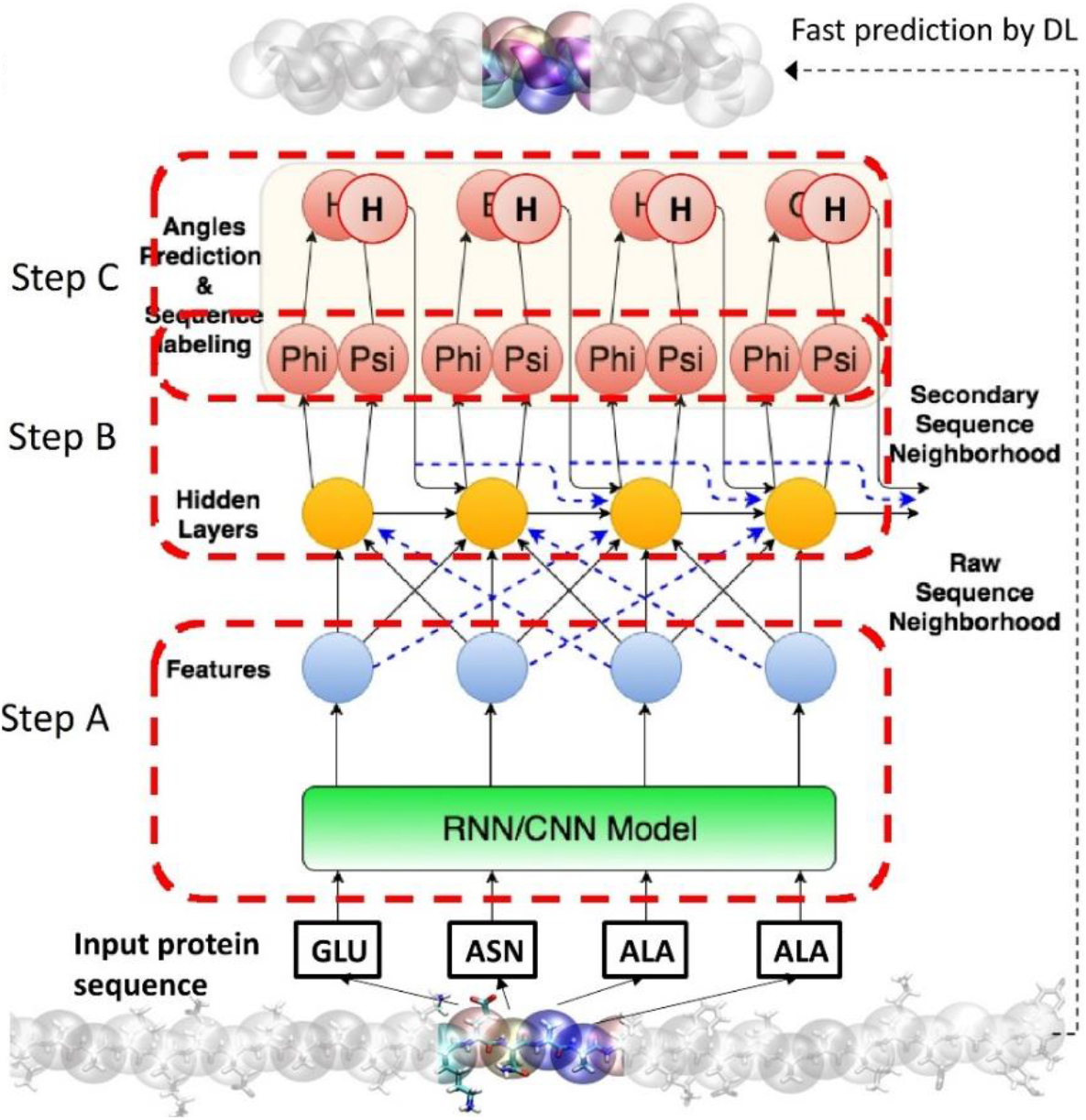

## Introduction

The development of rational techniques to discover new proteins for use in variety of applications ranging from agriculture to biotechnology remains an outstanding materials design problem (Ebrahimi et al., 2015; Gagner et al., 2014; Gronau et al., 2012; Kim et al., 2015; Selberg et al., 2018; Zhong et al., 2014). In fact, proteins represent the key construction materials of the living world, and offer enormous diversity in function, and hence a powerful platform for potential for use in bioengineering, medicine and materials science. Among several universally found secondary structures, alpha-helices (AHs) are a universal motif found in many protein materials. These protein domains play a crucial role in the signaling and deformation behavior of cytoskeletal protein networks in cells (e.g. intermediate filaments, as well as actin) (Gan et al., 2016; Herrmann and Aebi, 2004; Rowat et al., 2008), and dictate the mechanical elasticity of hair, hoof, feather and many other important structural protein materials (e.g., keratin) (Windoffer et al., 2011). Several silk proteins also possess helical structures, which impart toughness and antimicrobial functions to the final material (Sutherland et al., 2011, 2018; Weisman et al., 2010). AHs are also the most common structural motif in cellular membrane proteins, which are responsible for transport of matter, cell recognition, docking and signal transduction (Von Heijne, 2006; Selberg et al., 2018). In these materials, nanostructured AH based protein domains universally define their nanoscale architecture and biological functions. Despite the relatively simple geometry and well-known mechanical functions of alpha-helical domains, fast predictions of their assembly via non-helical domains to form a 3D folded structure is crucial to identify biological functions of a protein (e.g., binding, docking, assembly into higher-order structures, or specific biological properties such as antimicrobial).

Although the Protein Data Bank (PDB) (Tama et al., 2000) provides a rich resource of folded protein sequences and their all-atom 3D geometries (~120,000 protein structures to date), this database only includes a tiny portion of all proteins known to exist. Most proteins, however, are only known by their sequence and a limited set of associated functions (such as the 147,413,762 protein sequences given in uniprot.org (Consortium, 2019)), and their high-resolution protein structure remains unknown. Indeed, it is difficult to identify the complex structure of a protein from a pure experiment, which requires advanced tools including Nuclear Magnetic Resonance, X-Ray Diffraction or Cryo-electron microscopy, as well as protein crystal samples. Many proteins cannot be investigated in that way and hence, their full 3D structure, full set of functions (including roles in disease etiology or as platform for biomaterials) remains elusive. Structural homology helps to guess the 3D structure base on sequence similarity between the query protein and proteins with known structures used as templates (Zhang, 2008). However, the methods are often limited to analyzing small proteins due to the high computational cost, and they are unable to predict structures for which no existing protein templates or co-evolution information. Moreover, since functional proteins heavily occupy the PDB, the prediction becomes less relevant or absent for structural proteins sequences for designing materials targeting at advanced mechanical functions (Daga et al., 2010).

The protein dynamics revealed by atomistic simulation in an accurate solvent model condition can provide an accurate *ab initio* description of how a protein changes its conformation toward the state with lower free energy (Daga et al., 2010). It has been demonstrated that the 3D folded structure of a short protein sequence can be obtained from a computational simulation of protein folding directly from the sequence information (Conchúir et al., 2015; Cooper et al., 2010). However, the final equilibrated structure greatly depends on the initial conformation, as the structure can easily be trapped at a local energy minimum, while the global energy minimum can only be reached by crossing energy barriers, which are exceedingly rare transition events that must happen during a classical molecular dynamics (MD) simulations (Voelz et al., 2010). A typical protein of ~100 amino acids could require few seconds to fold (Naganathan and Muñoz, 2005). As classical MD computes the interactions and motions of a large number of atoms stepwise and as each time increment must be on the order of 1~2 fs (Naganathan and Muñoz, 2005), it would require 10^15^ computational integration operations for the full simulation of the folding trajectory, which goes beyond the capability of most supercomputers. Other methods such as the Replica Exchange Method (Sugita and Okamoto, 1999) effectively combine different simulation algorithms to accelerate the folding calculation compared with classical MD, but are still not fast enough to provide rapid results.

Artificial intelligence (AI), enabled by deep learning (DL) techniques, has demonstrated its advantage in solving sophisticated scientific problems that involve multiple physics interactions that are challenging to directly model or non-polynomial problems that require extremely large computational power that cannot be solved by brute force (Silver et al., 2016; Yu et al., 2019). Recent work has suggested that it may provide a feasible way to achieve fast prediction of protein structures by utilizing efficient algorithms of searching a high-dimensional parameter space for the most accurate prediction. In several materials-focused studies, such a data-driven material modeling for optimized mechanical properties of materials, it has shown its great potential in advancing conventional multiscale models in terms of efficiency and speed of predictions (Gu et al., 2018a, 2018b; Hanakata et al., 2018; Yu et al., 2019). Recently, deep learning techniques have shown progress in building an end-to-end differentiable model or directly predicting dihedral angles (AlQuraishi, 2019) or 3D structures of proteins (Evans, R.; Jumper, J.; Kirkpatrick, J.; Sifre, L.; Green, T.F.G; Qin, C.; Zidek, A.; Nelson, A.; Bridgland, A.; Penedones, H.; Petersen, S.; Simonyan, K.; Crossan, S.; Jones, D.T.; Silver, D.; Kavukcuoglu, K.; Hassabis, D.; Senior, 2019; Liu et al., 2018; Senior et al., 2018; Wang et al., 2018). However, all deep learning methods reported thus far still heavily rely on domain specific input features beside the primary amino-acid sequence, making it hard to generalize new protein sequences with no other known information about these input features. Moreover, the accuracy of these published works is validated within the PDB, but no application have been shown for innovative structural protein with unknown structure absent from the PDB, which will be essential for designing protein material from scratch.

In this paper, we present the design, training and validation of a Multi-scale Neighborhood-based Neural Network (MNNN) model for predicting the 3D all-atom structure, by exploiting the sequence neighborhood information from both the amino acid type of the secondary structure and its primary neighboring sequences. It is worth noting that the prediction takes only primary amino-acid sequence as inputs and makes structural prediction without any template or co-evolutional information. To accurately label the structure of each amino acid, we adopt K-means clustering to compute the possible number of secondary structure classes instead of directly applying the conventional eight-class categorization as commonly given by DSSP (Sreerama and Woody, 2000). Our experimental results show that our MNNN model can accurately predict whether a sequence can form an alpha-helical domain and its 3D all-atom structure with small prediction errors (see STAR Methods for details).

## Results

### Integrated Framework for Protein Design

We report a framework that integrates collective effort of MNNN, MD and experimental protein synthesis for alpha-helical protein designs, as shown in **Fig. 1**. Compared to experience-based trial-and-error for protein synthesis, this framework allows high-throughput *in silico* prediction of protein structures and related material functions that provide a rational basis for the design of *de novo* protein materials. One key driving force for design is to learn how the 3D protein structures are mapped to specific peptide sequences from all the known protein structures and to make fast prediction of the 3D structure of helical and non-helical domains from sequence. To achieve this goal, we download and analyze each of the 120,000 folded protein sequences and their all-atom 3D geometries by extracting the amino acid type and phi-psi dihedral angles of each amino acid, which quantitatively describes how the backbone atoms of the amino acid connects to its neighboring ones. It is noted that only phi-psi angles are considered in our model because other geometric parameters, such as omega, bond length and angle have constant values for alpha-helical structures and are taken these default values during prediction. We thus build a database to summarize the sequence information together with the phi-psi angle of each amino acid of the 120,000 folded protein structures (the data file is available for downloading). Our MNNN model learns how the phi-psi angles are mapped to specific peptide sequences and can then make predictions of the phi-psi angles for any given protein sequence. These predicted angles, combined with the default values of geometric parameters, are then used to build the coordinates of the backbone atoms of the proteins. The sequence information is thereafter used to decide the type of sidechain for each amino acid that connects to the center atom of the corresponding backbone atoms to build an all-atom 3D folded protein structure that can serve as the input geometry for further refinement and characterization of material functions through conventional MD simulations (see STAR Methods for details), such as protein’s thermal stability and diffusion coefficient at different temperatures. This multi-stage modeling process can effectively give the fast prediction of the 3D protein structure for any known or unknown protein sequences, which provides a helpful tool to sequence design.

**Figure 1:**
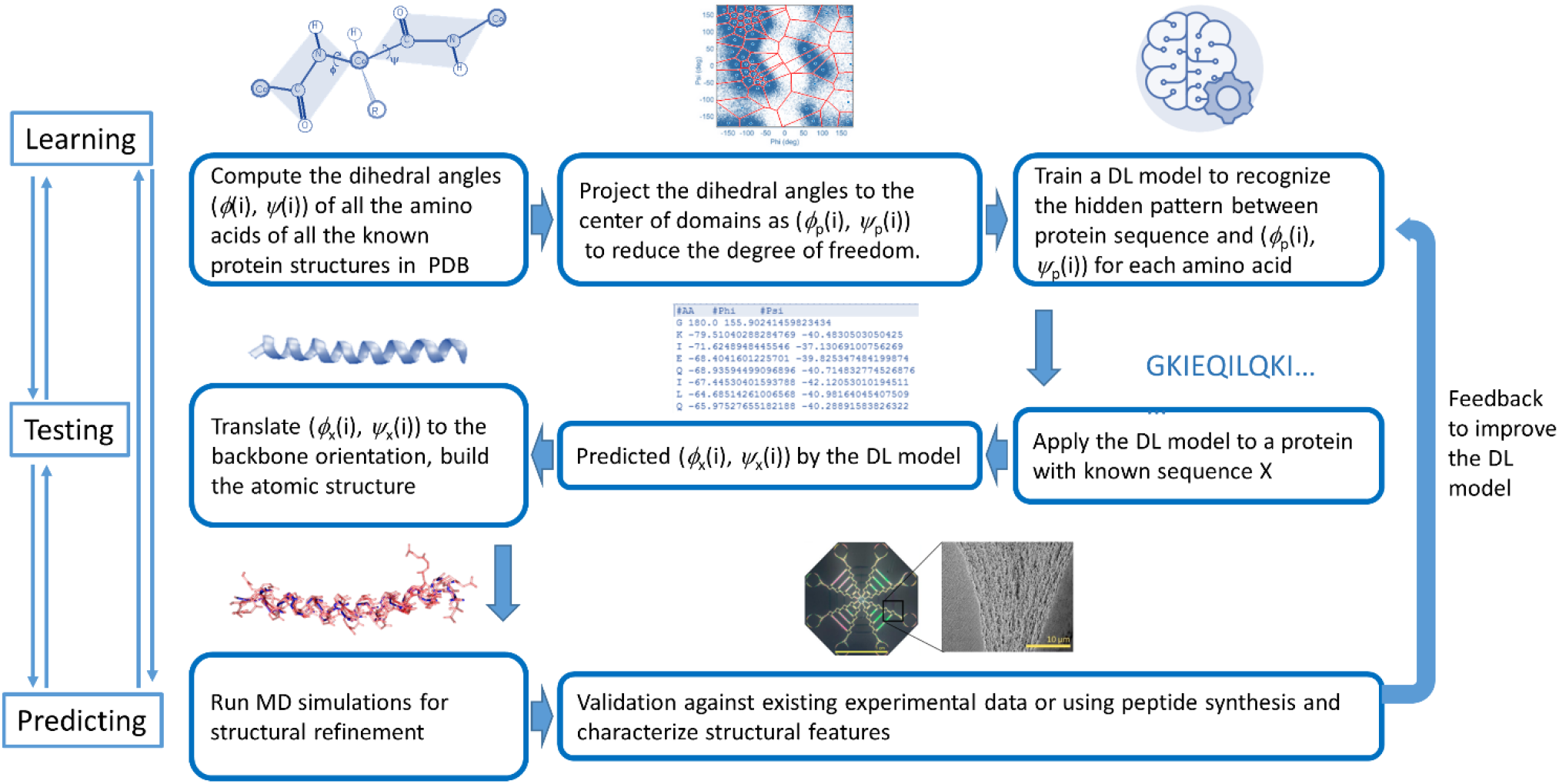
Overall flowchart of the algorithm reported in this paper. Taking the entire Protein Data Bank (composed of ~120,000 protein structures) as the training set, we extract the sequence information, along with the phi-psi angle information from each Protein Data Bank file for the high-resolution protein structure. We further label the dihedral angles by clustering according to the natural distribution of phi-psi angle to reduce the degree of freedom and use the structural labels, combined with the sequence information to train a MNNN model, which allows us to predict the phi-psi angle of any sequence and thus build the atomic structure together with the other intrinsic coordinate parameters. We run long-time MD simulations to quantify the acceleration of MNNN prediction and the stability of the structure given by MNNN result. The result is compared with experimental synthesis and characterization, which provide feedback to improve the quality of the MNNN model.

### Architecture of Multi-scale Neighborhood-based Neural Network for Dihedral Angle Prediction

Our deep learning regression model for predicting dihedral angles directly from the sequence neighborhood is based on a data-driven partition of design space of a protein structure, as shown in **Fig. 2**. The model is an end-to-end machine learning system that only requires raw protein sequences as data inputs and produces phi-psi angle prediction as outputs. To achieve the goal, we notice that the existing partition of the design space of protein structures (e.g., established methods such as DSSP (Kabsch and Sander, 1983)) only considers an eight-class human-engineered categories, which are too coarse to accurately characterize the diversity of natural structures. For example, taking a look at a typical Ramachandran plot, which summarizes the backbone dihedral phi-psi angles amino acid residues in different protein structure, a secondary structure as alpha-helix defined by DSSP may correspond to a wide range of phi value from −57° to −130° and psi value from −47° to −70°, while a single phi-psi value of the protein backbone will be crucial to precisely reconstruct the protein 3D structure. To address this challenge and quantify how one folded structure differ from others, we label each amino acid with a secondary structural information defined in a very different way from DSSP. Given the targeting number of categories, this data-driven approach computes the boundaries of the categories by considering the distribution of the phi-psi angles of all the known protein structures to minimize the within-cluster sum (**Fig. 2A**, see STAR Methods). We use clustering techniques such as K-means clustering on all PDB data with different cluster numbers and verify its degree of matching in comparison with the benchmark PDB structure, as shown in **Figs. 2A** **and** **B**. We find that when the class number is set to 256, the error between simulated structure with the benchmark result is reduced to a small level.

**Figure 2:**
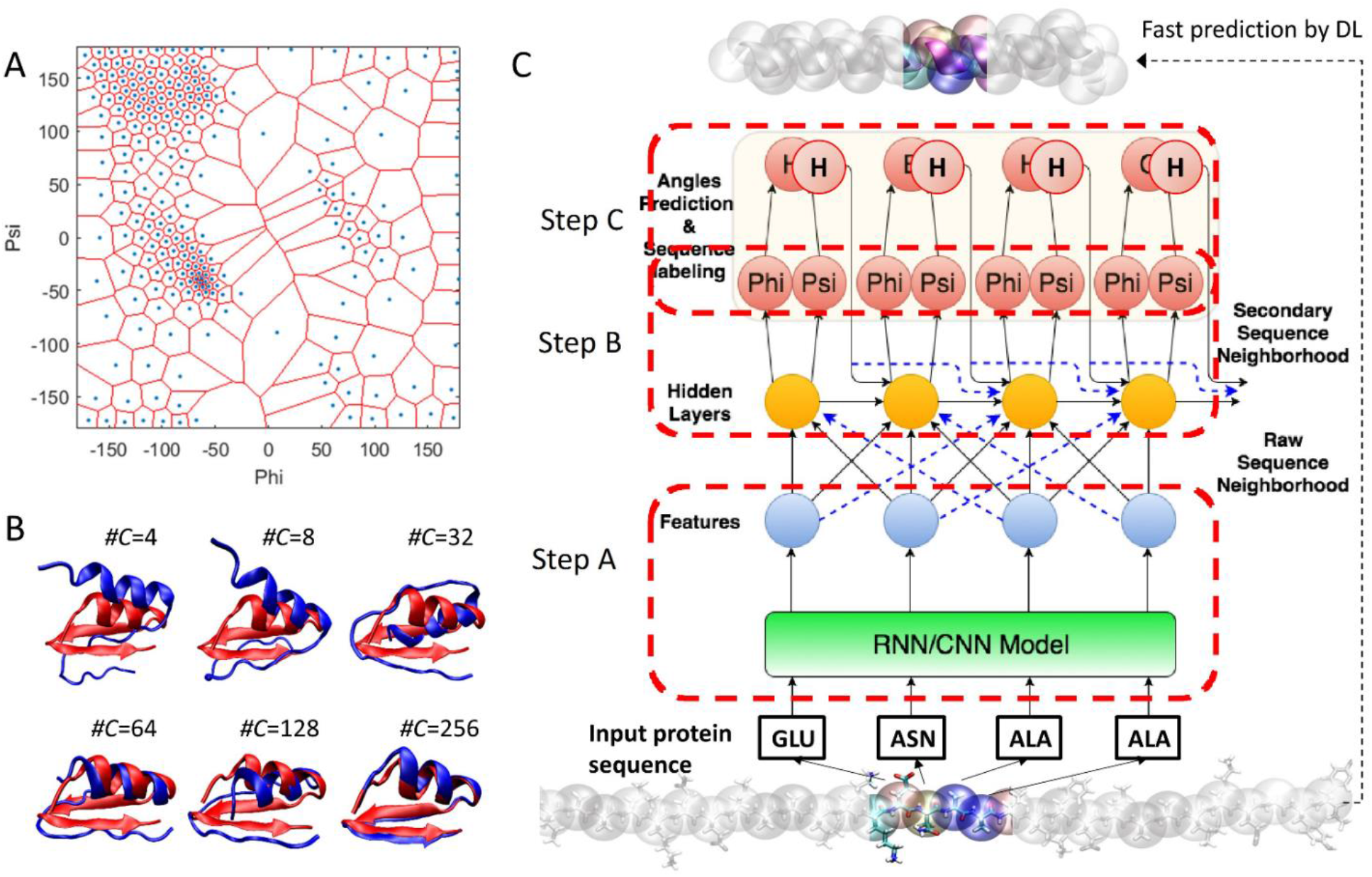
Strategy to reduce the design space of backbone conformations while achieving a high fidelity and overall architecture of our proposed MNNN model. A) We use a K means clustering algorithm to categorize all phi-psi angles in PDB into 256 clusters, which effectively reduces the infinite combination of phi-psi angles to the value of one of the cluster center (0..255). B) This panel shows the performance of using a different number of clusters in representing a protein structure (blue color, PDB ID: 1ACW), in comparison with the PDB structure (red). It is shown that for a 256 cluster choice, the error is reduced to a much smaller level with RMSD of 0.96 Å, comparing to RMSD of 7.3 Å, 6.44 Å, 6.6 Å, 3.1 Å and 4.3 Å for 4, 8, 32, 64 and 128 clusters, respectively. C) The architecture of our MNNN model (step A, B, and C) takes into account both information of raw sequence neighborhood and of secondary sequence neighborhood. In step A, we compute character embedding for each amino acid using any popular techniques and then refine these embeddings with sequence neighborhood information. In step B, we apply another LSTM layer to perform dihedral angles prediction based on refined character embeddings of amino acid sequence. In Step C, we use these two embeddings to further predict secondary structure characters as additional constraints based on predicted dihedral angles.

Since the secondary structure of neighboring amino acids will influence the subsequent secondary structure of the next amino acid, it is important to consider the neighboring K number of secondary structure prediction information when predicting dihedral angles of the next amino acid. When training a MNNN model (**Fig. 2C**), both data embeddings representing the raw amino acid sequence and their secondary structure information, as defined by our own, are incorporated to learn a refined embedding. This provides important structure information about the raw amino-acid sequence and thus serves as additional constraints for a MNNN model to achieve a better angle prediction. By considering both raw sequence and structural information in training, our model is able to learn the sequence-structure correlation among the subsequent continuous amino-acids. We then use the refined embeddings of the neighboring amino-acids for phi-psi dihedral angle predictions of a given sequence of amino acids.

### Benchmark for Prediction of Protein Structures

For structure predictions, we find that the MNNN model exhibits a significant advantage in speed and accuracy over the popular homology modeling and *ab initio* protein folding, such as all-atom MD simulations. For validation, we selected ~2.6 million short sequences (between 10 and 100 amino acids) with their structures known as alpha-helix or non-alpha-helix (by characterizing their structural files with DSSP (Kabsch and Sander, 1983)) for testing by MNNN (see STAR Methods for details, the facts and results of these testing sequences are available for downloading). Using this large number of testing cases, we find our prediction accuracy reaches 85.0%, which is much higher than the accuracy of many homology based secondary structure prediction (up to 80% (Daga et al., 2010)) and it is important to point out that the current method does not require significant sequence similarity (usually >30%) between query and structure for homology prediction (Daga et al., 2010). It is also higher than former learning methods (71.2%) (Wang et al., 2008) or predictions based on a genetic algorithm (75.1%) (Won et al., 2007), neural networks (78.1%) (Yao et al., 2008) or support vector machine (82.2%) (Duan et al., 2008). This is very important for design as the high accuracy and low constraint is crucial to make the tool efficient in identifying the distribution of helical structures within an unknown sequence.

In comparison to *ab initio* protein folding by molecular dynamics, the current method shows great advantage for predicting speed and accuracy. For benchmark, six well-characterized coiled-coil peptides (Burgess et al., 2015; Fletcher et al., 2012; Thomson et al., 2014; Zaccai et al., 2011) were chosen as benchmark proteins to test the efficiency and accuracy of our MNNN model. The backbone root-mean-square deviation (RMSD, see STAR Methods) of the MNNN-predicted structures, as the deviation of the coordinates all the four backbone atoms (N, CA, C, O) of each amino acid, is compared to the available structures in the PDB for the six peptides were computed. The outcomes, as summarized in **Table 1**, are compared against protein folding with classical MD simulations with implicit and explicit solvent models. It is shown that the largest error given by MNNN is merely 2.11 Å, which is much better than folding the peptide from a fully extended form in MD, either in implicit (12.9±4.2 times the error) or explicit (10.4±4.6 times the error) solvents. Most importantly, the time needed to obtain the predicted folded structure from MNNN is significantly smaller (less than six orders of magnitude) than classical MD simulations (with the least requirement of computing hardware). This outcome suggests that the MNNN model can offer a powerful way to predict protein structures.

**Table 1:**
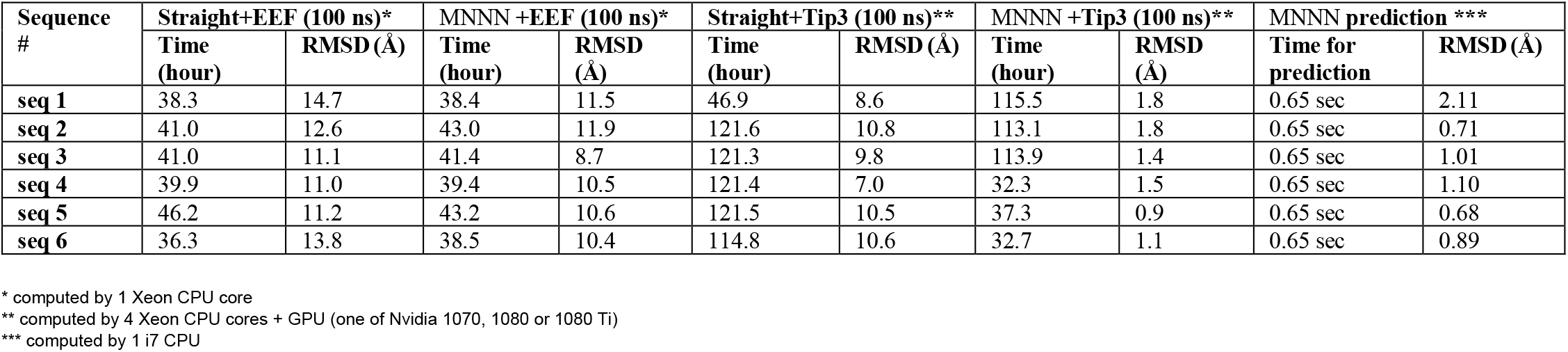
This table summarizes the computational efficiency for different simulation and prediction methods and their error (backbone RMSD as given by Eq. (1)), by comparing to the 3D structure as well as experimental results.

| Sequence # | Straight+EEF (100 ns)* |  | MNNN +EEF (100 ns)* |  | Straight+Tip3 (100 ns)** |  | MNNN +Tip3 (100 ns)** |  | MNNN prediction *** |  |
| --- | --- | --- | --- | --- | --- | --- | --- | --- | --- | --- |
|  | Time (hour) | RMSD (Å) | Time (hour) | RMSD (Å) | Time (hour) | RMSD (Å) | Time (hour) | RMSD (Å) | Time for prediction | RMSD (Å) |
| seq 1 | 38.3 | 14.7 | 38.4 | 11.5 | 46.9 | 8.6 | 115.5 | 1.8 | 0.65 sec | 2.11 |
| seq 2 | 41.0 | 12.6 | 43.0 | 11.9 | 121.6 | 10.8 | 113.1 | 1.8 | 0.65 sec | 0.71 |
| seq 3 | 41.0 | 11.1 | 41.4 | 8.7 | 121.3 | 9.8 | 113.9 | 1.4 | 0.65 sec | 1.01 |
| seq 4 | 39.9 | 11.0 | 39.4 | 10.5 | 121.4 | 7.0 | 32.3 | 1.5 | 0.65 sec | 1.10 |
| seq 5 | 46.2 | 11.2 | 43.2 | 10.6 | 121.5 | 10.5 | 37.3 | 0.9 | 0.65 sec | 0.68 |
| seq 6 | 36.3 | 13.8 | 38.5 | 10.4 | 114.8 | 10.6 | 32.7 | 1.1 | 0.65 sec | 0.89 |

\* computed by 1 Xeon CPU core
\*\* computed by 4 Xeon CPU cores + GPU (one of Nvidia 1070, 1080 or 1080 Ti)
\*\*\* computed by 1 i7 CPU

Moreover, it is also important to note that the structures given by the MNNN model not only agree well with available structures in the PDB, but also yields predictions with good overall thermal stability. As summarized in Table 1 and shown in **Fig. 3**, the structure predicted by the MNNN model only appear to have a very small thermal fluctuations in all-atom MD simulation (with backbone RMSD<1.8 Å). It is also noted that the implicit solvent model (much less computationally expensive than explicit solvent models), no matter whether they start from the fully extended form or from the structure given by the MNNN model, does not yield good predictions for either folding or structure refinement purposes.

**Figure 3:**
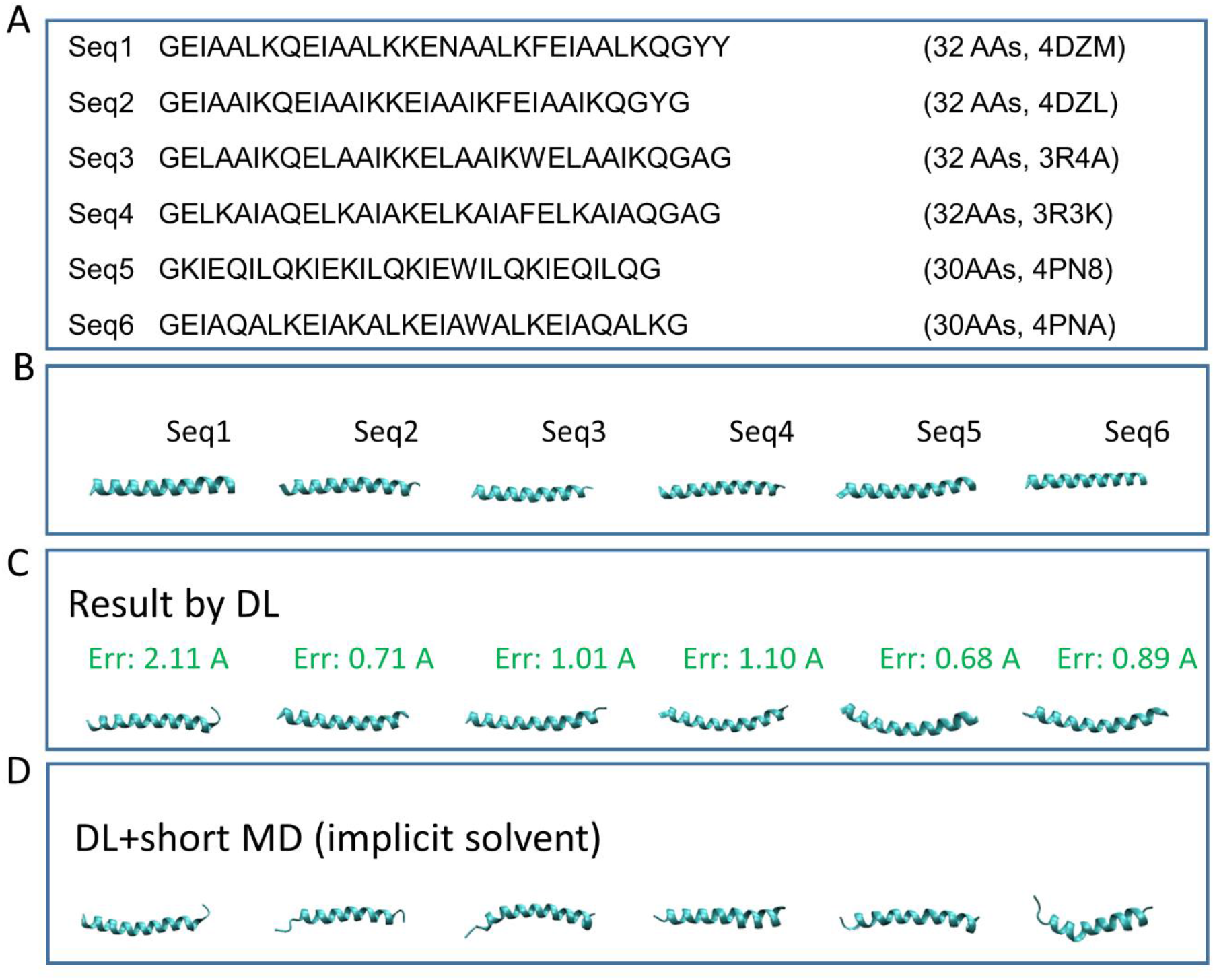
Summary of all sequences investigated for benchmarking, and a brief summary of the prediction results. A) The six sequences and their corresponding structure id in PDB. B) The snapshots of the protein structures as obtained from the PDB. C) The result of MNNN and the backbone RMSD value from PDB structures given in panel B.

### New Protein Design and Structural Validation

Besides the six small proteins whose 3D structures are already resolved, we test the accuracy and efficiency of our MNNN algorithm in protein folding prediction on sequences whose molecular structure is unknown. To this end, a peptide of 28 amino acid length (named AmelF3_+1) extracted from the coiled-coil domain of the Apis mellifera silk protein (AmelF3) (Sutherland et al., 2011) was chosen (**Fig. 5A**). The results of the structure of AmelF3_+1 as obtained by the MNNN model developed in this study is compared with structure homology prediction tools including Optimized Protein Fold Recognition (ORION) (Ghouzam et al., 2016) and Iterative Threading Assembly Refinement (I-TASSER) (Zhang, 2008), as shown in **Fig. 5B**. ORION is a sensitive method based on a profile-profile approach that relies on a good description of the local protein structure to boost distant protein structure predictions, while I-TASSER features a hierarchical approach for protein structure and function prediction that identifies structural templates from the PDB, with full-length atomic models constructed by iterative template-based fragment assembly simulations. The prediction by our MNNN model is an alpha-helix, which overall agrees with the results of the other two methods, both of which also predict alpha-helical proteins. We use the three predicted structures and compare their dynamic behaviors in 100 ns MD simulation in explicit solvent (**Fig. 5B**). It is found that the structure predicted by the MNNN model is more thermodynamically stable than the other two, particularly significantly better than the I-TASSER model, which almost unfolded during the middle of the simulation run. ORION is limited by template availability, and the prediction of the highest scoring structure may not be of the same length as the targeted sequence and one may need to balance between structure integrity and accuracy.

**Figure 4:**
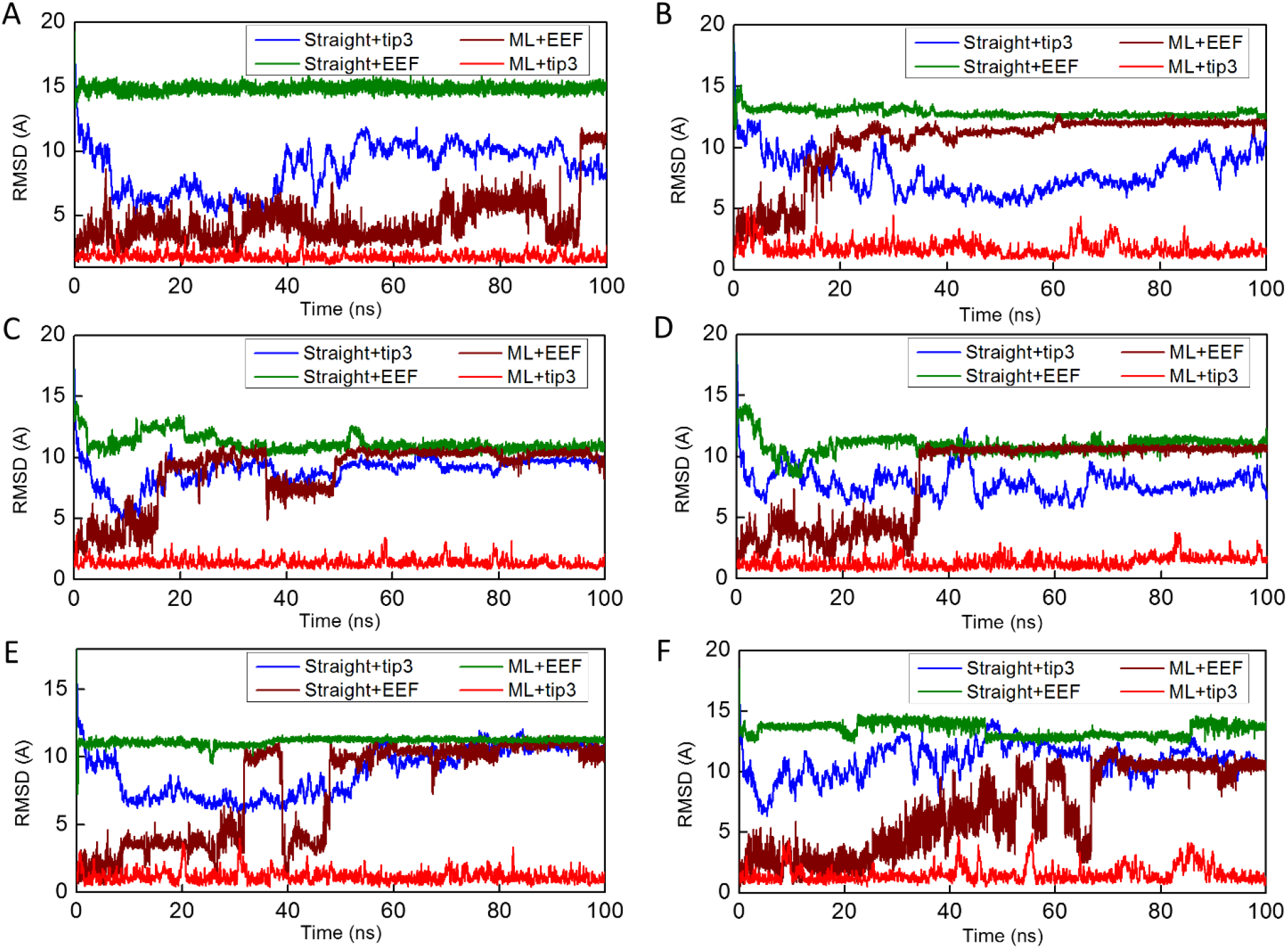
Benchmark results of all six sequences as summarized in Fig. 4. Each of them include different starting conformations (arbitrary straight chain and result obtained by MNNN) and different force fields and solvent models (Tip3P explicit solvent and EEF1 implicit solvent), with panels (A), (B), (C), (D), (E) and (F) for protein 4DZM, 4DZL, 3R4A, 3R3K, 4PN8 and 4PNA, respectively.

**Figure 5:**
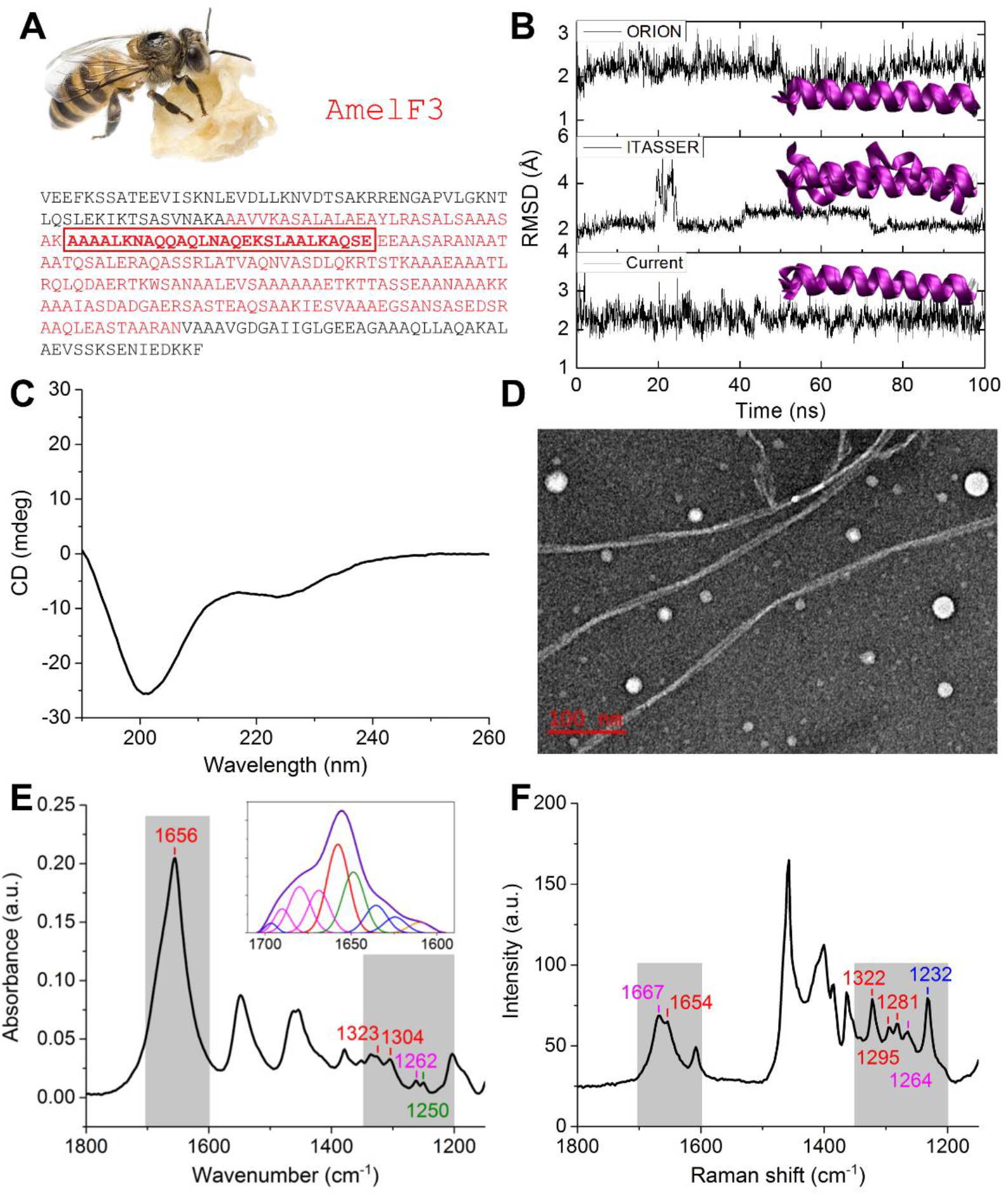
Synthesis and characterization of a *de novo* peptide sequence that is not part of the Protein Data Bank. A) Amino acid sequence of honeybee silk protein AmelF3, with the coiled-coil domain highlighted in red and the peptide (AmelF3_+1) sequence shown in **bold**. B) Results of 100 ns MD simulation starting from the predictions given by ORION, I-TASSER and the MNNN model. Snapshots taken every 20 ns of each simulation are overlaid for comparison. C) CD spectrum of AmelF3_+1 in deionized water. D) Representative transmission electron micrograph of the AmelF3_+1 peptide, depicting nanofiber formation. E) FTIR spectrum of AmelF3_+1 film, with alpha-helical peaks labeled in red, turns in magenta, and random coils in olive. The inset shows Fourier self-deconvolution and secondary structure peak fitting of the Amide I band. F) Raman spectrum of AmelF3_+1 film, with the peaks labeled by the same color notation for secondary structures as used in the FTIR spectrum, blue represents beta-sheet.

To validate the computational results, the peptide AmelF3_+1 was synthesized and characterized experimentally. The circular dichorism (CD) spectrum of AmelF3_+1 (**Fig. 5C**) shows a major combination of alpha-helical, beta-turns and random coils conformations (Greenfield, 2007; Kelly et al., 2005), with the relative contents being 57%, 24% and 14%, respectively, as estimated by the CONTINLL program (Greenfield, 2007; Sreerama and Woody, 2000). Moreover, potential peptide assembly into higher-order structures was captured by TEM, as shown in **Fig. 5D**, from which we can see that the peptide assembles into either nanofibers of around 10 nm in width and several microns in length or nanoparticles of diameters ranging from 10-35 nm. The secondary structure analysis was also independently confirmed by ATR-FTIR and Raman spectroscopy, the spectra of which are complementary in the Amide I and III bands, with well-established peak assignments for different secondary structures (Hu et al., 2006, 2008; Woodhead et al., 2016). From the FTIR spectrum of AmelF3_+1 (**Fig. 5E**), a major peak at 1656 cm-1 in the Amide I region along with the two peaks at 1323 and 1304 cm-1 in the Amide III region indicate predominant alpha-helical conformations of the peptide. The Raman spectrum of AmelF3_+1 (**Fig. 5F**) gives similar structural information, with two more hidden alpha-helical peaks (*i.e.*, 1295 cm-1 and 1281 cm-1) clearly seen. It is anticipated that with the correct buffer condition, more stable alpha-helical conformation of the AmelF3_+1 peptide can be achieved.

## Discussion

In this paper we reported a MNNN model that requires no templates, co-evolution information or structural biological knowledge to accurately and rapidly predict the structure of *de novo* proteins directly from the primary protein sequence. We show it can reliably (of 85% prediction accuracy) distinguish helical domain from non-helix domain without requiring significant similarity between query and structure, as what is needed for homology predictions. We evaluate the quality of the predicted 3D structures and it shows a maximum error of merely 2.1 Å for all the tested sample sequences. We further investigate its accuracy and application by predicting the 3D helical structure of honeybee silk protein that is not known for its protein structure and our prediction agree with the protein synthesis and the results of experimental characterizations very well.

MNNN is implemented as a freely available and flexible tool that requires minimum computational resource in structure prediction. The prediction of the structures of 2.6 million short sequences takes only a few hours on a laptop, making its implementation to a webserver quite feasible for public access and evaluation of the prediction reliability broadly. Since the MNNN learns from all known protein structures, its prediction accuracy can be further improved by incorporating new structure to refine the models as new proteins being discovered daily. At the same time, its predicting speed will not be affected as the new data will only be inferred using the parameters in the neural network but will not be used as templates during the prediction.

The MNNN model introduced here represents a significant advancement over previous methods in predicting protein structures as shown in Table 2. Compared to classical ML methods, our method MNNN does not requires template (explicit s2s maps) or co-evolutional information to predict 3D structure. Another huge advantage of MNNN is that it can perform prediction in an ultrafast speed. Compared to very recent DL methods (AlphaFold and DGN), our method is only method that explore the neighborhood information of both raw sequence and the secondary structure during training and perform prediction solely relying on raw sequence. For instance, AlphaFold cannot directly take only raw sequence and had to use co-evolving residues information to perform prediction, while DGN needs to take both PSSMs and raw sequence to perform prediction and it cannot leverage secondary structure information yet.

**Table 2:** This table summarizes the various information different AI methods need and potential advantages and disadvantages of each type of methods. S2s stands for sequence to structure.

| *AI method* | Use<br>explicit<br>s2s maps | Use co-evolving<br>residues | de novo<br>protein<br>design | Handle Minor<br>sequence<br>mutations or<br>indels | Use<br>PSSMs | Use<br>secondary<br>structure | Computational<br>cost |
| --- | --- | --- | --- | --- | --- | --- | --- |
| *Classical method I* | Yes | No | Yes | Yes | Yes | No | Very slow |
| *Classical method II* | No | Yes | No | No | Yes | No | Very slow |
| *AlphaFold* | No | Yes | No | No | No | No | Fast |
| *DGN* | No | No | Yes | Yes | Yes | No | Very Fast |
| *MNNN* | No | No | Yes | Yes | No | Yes | Very Fast |

This AI based approach to design new proteins opens the door to generative methods that can complement conventional protein sequence design methods. For example, it can be used to design a long sequence that will yield the desired distribution of alternative helical and coil domains to assembly into functional channels for delivery of matters. Future work could expand the method to include other secondary structures, and achieve a broader, more comprehensive structure prediction capacity.

## STAR*Methods

### Data Preparation for Training and Translating Predictions to 3D All-Atom Protein Structures

We develop a custom script to obtain the phi-psi angle and sequence information of each of 126,732 natural protein structures composed of only standard amino acids that are currently available in the PDB. We wrote our own bash script that allows using DSSP to compute the phi-psi angles of the backbone by reading the 3D structure files that are automatically downloaded from PDB and using open source Unix software to build a highly structuralized database for training (**Fig. 1**). We develop a post-process script to take the (phi, psi) angle as predicted by MNNN to combine with the rest of the geometric parameters given by the intrinsic coordinates within the CHARMM force field to build the all-atom protein structure, which allow to run energy minimization and molecular dynamics to validate the thermal stability of the protein structure.

### Neural Network Training and Validation

The MNNN model predicts dihedral angles directly from sequence neighborhood, by taking into account both raw sequence neighborhood and secondary sequence neighborhood. Our model consists of three main steps:

A. We compute character embedding for each amino acid (can be done using a variety of popular techniques) and then refine these embeddings with sequence neighborhood information. For instance, one can use long short-term memory (LSTM)(Hochreiter and Schmidhuber, 1997) or bidirectional LSTM (Bi-LSTM) (Graves and Schmidhuber, 2005) to compute character embeddings. One can also use other character embedding techniques such as fasttext (Joulin et al., 2016), Glove (Pennington et al., 2014) or Transformer (Vaswani et al.) to compute character embeddings. In this paper, we choose to first use a Bi-LSTM layer to compute character embeddings of primary amino-acid. Then we apply a convolutional neural network (CNN) (Krizhevsky et al., 2012) layer to take into account sequence neighborhood information based on computed character embeddings of each amino acid.
B. Based on refined character embeddings of amino acid sequence, we apply another LSTM layer to perform dihedral angles prediction. For the first amino acid, it uses only its own hidden representation to predict dihedral angles. For the second and subsequent amino acids, we refine the embedding of current animo acids using the previous K number of embeddings of predicted secondary structure character and then predict dihedral angles.
C. Once dihedral angles are predicted, we use these two embeddings to further predict secondary structure characters as additional constraints. The existing partition of the design space of protein structures is divided into an eight-class human-engineered categorization. However, this choice is ad-hoc and there are large sets of structural features that are not well covered by this categorization. Therefore, we propose to use a data-driven approach (clustering techniques such as K-means clustering) to compute the possible number of secondary structure classifications. We use a K-means clustering algorithm to categorize all phi-psi angles in PDB into 256 clusters, which effectively reduces the infinite combination of phi-psi angles to the value of one of the cluster center as shown in Figure 2 (A) and (B). In other word, instead of using conventional 8-class categorization of secondary structure, we leverage data-driven approach to use new 256-class categorization of secondary structure during the data preprocessing. During training, we use our new 256-class categorization of secondary structure for labeling the structure of each amino acid and train our model as shown in Figure 2 (C).

Our overall loss function for our MNNN model consists of three parts: i) a loss function with RMSE of the predicted Phi angle and the real ones; ii) a loss function with RMSE of the predicted Psi angle and the real ones; iii) a loss function for matching the predicted secondary structure class with real classes (from data-driven approach). Note that the angles are represented by trigonometric functions (*i.e*., an angle is converted to a pair of its sine and cosine values for numerical stability during model training). We then split the PDB data into train/validation/test datasets and train the model with train/validation data and test our models with test data. We further compute the L1 norm of dihedral angles of any target sequence within average L1-norm errors for the phi-psi angle pair. Throughout our experiment, we set the neighborhood parameter in our MNNN model to be 21. On the test dataset, we obtain average L1 errors of 22 degrees and 37 degrees for the Phi-Psi angles respectively. The training error is similar to the test error.

### Prediction accuracy

We run predictions for a large-scale collection of sequences to carefully compute the accuracy in helix prediction. We selected 2.6 million short sequences with length between 10 and 100 amino acids from PDB with experimentally known structures. It is noted that this sample size is three orders of magnitude larger than people used to validate their prediction algorithm (Duan et al., 2008; Wang et al., 2008; Won et al., 2007; Yao et al., 2008), thanks to the high efficiency of our learning model. About half of them are sequences that make pure alpha helix domain (Qin et al., 2013), and the other half are totally non-helical domain. We use our MNNN model to predict the phi-psi angle, give their 3D structures and then use DSSP (Kabsch and Sander, 1983) to get the secondary structure for each amino acid within each sequence as either helix or non-helix structure. Thereafter, we compare the result with the experimental measurement by omitting the first and last amino acids as their secondary structures cannot be well defined with one dangling end, and obtained *N*_hh_=13,277,400, as the number of amino acids experimentally measured as helix and predicted as helix by MNNN, *N*_hn_=2,877,342, as the number of amino acid experimentally measured as helix and predicted as non-helix by MNNN, *N*_nh_=3,911,776, as the number of amino acid experimentally measured as non-helix and predicted as helix by MNNN, and *N*_nn_=25,250,846, as the number of amino acid experimentally measured as non-helix and predicted as non-helix by MNNN. We compute the accuracy of the prediction by using (*N*_hh_+ *N*_nn_)/(*N*_hh_+ *N*_hn_+ *N*_nh_+*N*_nn_)=85.0% which gives the prediction accuracy of our MNNN model.

### Molecular modeling

We use two force fields (CHARMM19 with implicit solvent (Lazaridis and Karplus, 1997), and CHARMM27 with explicit solvent (MacKerell et al., 1998)), with each of them starting from two different extreme configurations (full extended chain, and a structure with phi-psi angles predicted from MNNN), to perform individual long-time simulations (100 ns) for each of the protein sequence. During the simulation, the change of the molecular conformation is benchmarked by quantitatively comparing with the corresponding protein structure within the PDB.

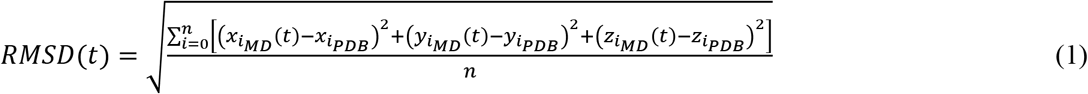

Where *n* is the number of all the backbone atoms of each amino acid of the peptide (N, C, CA, O) and 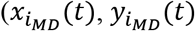 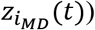 is the Cartesian coordinates of the backbone atoms given by the MD simulation at time *t*, while 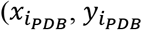 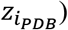 is the Cartesian coordinates of the backbone atoms of the protein structure within the PDB. In the CHARMM model, the mathematical formulation for the empirical energy function has the form:

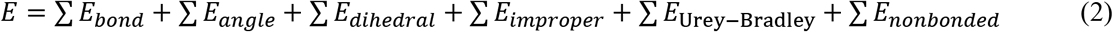

Each energy term is given by *E*_*bond*_ = *K*_*ij*_(*r* − *r*_0_)^2^ is the bond term that defines how two covalently bonded atoms interact in the stretching direction, *E*_*angle*_ = *K*_*ijk*_(*θ* − *θ*_0_)^2^ is the angle term that defines how the angel among three covalently bonded atoms with one central atom changes under external force, *E*_*dihedral*_ = *K*_*ijkl*_[1 + cos(*nϕ* − *δ*)] is the dihedral term that defines how the dihedral angel among four covalently bonded atoms with one central bond changes under external force, *E*_*improper*_ = *K*_*ijkl*_(*ω* − *ω*_0_)^2^ is the improper angle term that defines how the improper angle among four covalently bonded atoms with one central atom changes under external force, *E*_Urey-Bradley_ = *K*_*u*_(*u* − *u*_0_)^2^ is the Urey Bradley term that accounts for angle bending and 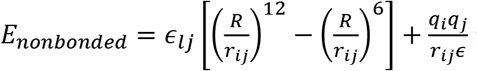 is the nonbonded term that accounts for the van Der Waals (VDW) energy and electrostatic energy. In all-atom force fields, water molecules generally can be treated either explicitly or implicitly for MD simulations.

We use the CHARMM19 all-atom force field to model the atomic interactions for the straight chain and the MNNN predicted model. The solvent effect for this force field is generally considered by using the implicit Gaussian model (EEF1) for the water solvent(Lazaridis and Karplus, 1997). The use of the implicit solvent model has advantages to accelerate the sampling speed of molecular configurations. We use the CHARMM c37b1 package to run the simulation for energy minimization and structural equilibration. Because there is no explicit water or pressure control, we do not apply any constraint to ensure the simulation stability. The time step used for implicit solvent simulations is 1 fs.

Starting from the initial geometry built using the backbone dihedrals (with (phi, psi)=(180°, 180°) for each amino acid for a straight chain, and (phi, psi) defined by MNNN for another model), combining with the rest of the geometric parameters given by the intrinsic coordinates within the CHARMM force field, we follow the following protocol to equilibrate the structure: 1) Energy minimization (2,000 Steepest Descent steps followed by 2,000 Adopted Basis Newton-Raphson steps); 2) Equilibration runs for 50 ps (NVT ensemble with Nose-Hoover temperature control), where the temperature rises linearly from 240 K (beginning) to 300 K (end); 3) Equilibration runs for 100 ns (NVT ensemble with Nose-Hoover temperature control), where the temperature stays at 300 K. We record the coordinates for each 10 ps, compare with the PDB structure by backbone RMSD to measure how far is the folded structure away from the PDB structure.

Besides implicit solvent, we use the CHARMM27 force field implemented by the explicit TIP3P water model and run simulation with NAMD package v2.13(Nelson et al., 1996) with the support of Graphical Processing Units (GPUs), which greatly outperforms Central Processing Unit (CPU) performance. Our model starts from the equilibrated structure obtained by using the implicit solvent model described above. All simulations run in a NPT ensemble under a constant temperature (300 K) and constant pressure (1 atmosphere) controlled by Langevin thermostat and barostat. The simulation time step is 2 fs with rigid bonds model for all the covalent bonds between hydrogen atoms and other heavy atoms. We use particle mesh ewald (PME) function with a grid width <1 Å to calculate the electronic interaction because and it is an efficient method to accurately include all the long-distance electrostatics interactions. A cutoff length of 10 Å is applied to the van der Waals interactions and to switch the electrostatic forces from short range to long-range forces.

The initial protein structure for explicit solvent simulation is built the same way as the implicit models. We use Visual Molecular Dynamics (VMD) (Humphrey et al., 1996) to add a solvent box around the protein structure with water at a distance of least 10 Å from the protein structure. The net charge of the system is set to zero by adding NaCl of overall concentration of 0.1 Mol/L, and each ion is initially randomly placed in the solvent box with the actual ratio of ions adjusted to neutralize the system. We follow the following protocol to equilibrate the structure: 1) Energy minimization (10,000 conjugate gradient steps); 2) Equilibration runs for 100 ns (NPT ensemble), where the temperature stays at 300 K. We record the coordinates for each 10 ps, compare with the PDB structure by computing backbone RMSD to measure how far is the folded structure away from the PDB structure.

### Peptide synthesis

The peptide used in this study was synthesized by GenScript (Piscataway, NJ), with free N- and C-termini. Peptides were synthetized using standard Fluorenylmethyloxycarbonyl (Fmoc)-based solid-phase peptide synthesis (SPPS) and purified by reverse-phase high-performance liquid chromatography (RP-HPLC) to a purity of 95% or higher.

### Circular Dichroism (CD) Spectroscopy

Circular Dichroism (CD) spectra were recorded from 190 to 260 nm using a JASCO J-1500 spectrometer, with each spectrum averaged from three consecutive scans, the wavelength step being 0.5nm and the scan rate being 50 nm/min. Samples of 1mg/ml in deionized water were measured in a 0.1mm path length quartz cuvette (Starna Cells, Inc.). Secondary structure estimation was performed using the CONTINLL program with a reference set of 48 soluble proteins.

### Fourier-transform Infrared Spectroscopy (FTIR)

ATR-FTIR measurements were performed on a Nicolet 6700 FT-IR spectrometer (Thermo Scientific) equipped with a liquid-nitrogen-cooled microscope. Spectra were collected in reflection mode with ATR correction using a germanium crystal. Each spectrum was collected from 4000 to 650 cm^−1^ with a resolution of 4 cm^−1^ and 64 scans. The relative fractions of different secondary structures were determined by Fourier self-deconvolution (FSD) of the Amide I band (1705-1595 cm^−1^) and Gaussian curve-fitting of the deconvoluted spectra using Origin.

### Raman Spectroscopy

Raman spectroscopy were performed using a Renishaw Invia Reflex Raman confocal microscope equipped with a 1” CCD array detector (1024 × 256 pixels), a 532 nm laser and a 100× objective. Each spectrum was collected in the 101 - 2736 cm^−1^ range and as an accumulation of 20 scans to increase the signal-to-noise ratio (3 seconds exposure per scan).

### Transmission Electron Microscopy (TEM) imaging

Transmission electron micrographs were recorded using a Tecnai G^2^ Spirit TWIN (LaB6 filament, 120 kV) equipped with a Gatan CCD camera. Continuous-film carbon-coated copper grids (Ted Pella, CA) were glow discharged and used for negatively stained samples. Briefly, 5 μL peptide samples of 1 mg/ml in deionized water were pipetted onto the grid, wicked off after 2 minutes, washed with water and then stained with 5 μL 2% uranyl acetate for 1 minute before being wicked off. The grid was then left to dry before imaging. Dimensional measurements of the peptide assemblies were performed on ten different micrographs using DigitalMicrograph (Gatan Inc.).

## Acknowledgements

This research was supported by the IBM-MIT AI lab. The authors acknowledge the Center for Materials Science and Engineering (CMSE) and Biophysical Instrumentation Facility (BIF) at MIT for access to the structural characterization and imaging instruments.

## Author contributions

The research was designed by MJB, BM, LW and PYC with input from all co-authors. Molecular modeling and analysis was conducted by ZQ and MJB. The AI work was led by LW, SH, TM and PYC. The paper was written with input from all co-authors. All co-authors approved this submission.

## Data and Code Availability Statement

Sequence and the phi-psi angle of ~120,000 protein structures: https://www.dropbox.com/s/j5hk6yi9xsnc6u8/data_phi_psi_6_1.dat?dl=0

Experimental and predicting results of the 1.3 million test protein sequences as pure helical domain: https://www.dropbox.com/s/9zll9ile0ssndnw/pure_helix_structure_AI_all.dat?dl=0

Experimental and predicting results of the 1.2 million test protein sequences as pure non-helical domain: https://www.dropbox.com/s/ari0g8qmdmmi8gw/non_helix_structure_AI_all.dat?dl=0

Github link of MNNN codes for prediction of phi-psi angle of any protein sequence: https://github.com/IBM/mnnn Other Data and materials related to this paper may be requested from the authors.

